# Bayesian Modeling of Epigenetic Variation in Multiple Human Cell Types

**DOI:** 10.1101/018028

**Authors:** Yu Zhang, Feng Yue, Ross C Hardison

## Abstract

With massive amount of sequencing data generated for many epigenetic features in a variety of cell and tissue types, the chief challenges are to build effective and quantitative models explaining how the dynamics in multiple epigenomes lead to differential gene expression and diverse phenotypes. We developed a unified Bayesian framework for jointly annotating multiple epigenomes and detecting differential regulation among multiple tissues and cell types over regions of varying sizes. Our method, called IDEAS (*i*ntegrative and *d*iscriminative *e*pigenome *a*nnotation *s*ystem), achieves superior power and accuracy over existing methods by modeling both position and cell type specific regulatory activities. Using 84 ENCODE epigenetic data sets in 6 cell types, we identified epigenetic variation of different sizes that are strongly associated with differential gene expression. The detected regions are significantly enriched in genetic variants associated with complex phenotypes. Our results yielded much stronger enrichment scores than achievable by existing approaches, and the enriched phenotypes are highly relevant to the corresponding cell types. IDEAS is a powerful statistical tool for integrative annotation of regulatory elements and detection of multivariate epigenetic variation in many tissues and cell types, which could be of important utility in elucidating the interplay between genetic variants, gene regulation and diseases.

The states generated by IDEAS can be visualized or downloaded from the “Regulation” section of http://main.genome-browser.bx.psu.edu/

## Introduction

An essential problem in molecular biology is to understand gene regulation and its impact on phenotypic diversity. Applications of advanced biotechnologies^1-3^ in existing large consortia^4-9^ and many individual laboratories are generating massive data sets for many genome-wide features. Data acquisition is no longer the major barrier to understanding human epigenome, mechanisms of gene regulation and its role in complex disease. The chief challenges are to build effective and quantitative models explaining how the dynamics of epigenomes relates to gene expression changes and phenotypic diversity. Interpretation and modeling must be resolved both along the genome and across different cell/tissue types. Key regulators functioning in specific cell/tissue types can be revealed by contrasting epigenetic signals across samples, along with their impacts on gene expression^5^. New hypotheses about the dynamics of the regulatory regimes during differentiation can consequently be derived and tested to illuminate previously intractable issues in the genetics of disease susceptibility^10,11^.

Genetic variants for human complex diseases are significantly enriched in regions exhibiting epigenetic cell type-specificity^11,12^. While statistical endeavors have been conducted to identify cell type-specific epigenetic marks and pinpoint their locations^12-15^, there is a lack of generalized and rigorous methods to powerfully identify cell type-specific regions incorporating multiple epigenetic marks in many epigenomes jointly. A state-of-the-art approach for characterizing multivariate epigenetic marks is via hidden-Markov-model (HMM) enabled genome segmentation^16,17^, which translates the high-dimensional raw data into a set of comparable and interpretable states for segments of the genome exhibiting unique patterns of chromatin marks. The inferred epigenetic states have been proven useful, with experimentally confirmed functionality, as a convenient tool for studying gene regulation and their implications on phenotypes^5^. Two state-of-the-art algorithms are chromHMM^16^ and Segway^17^. ChromHMM is a multivariate hidden Markov model that interprets the presence or absence of chromatin marks over siding windows of fixed sizes. Segway uses a dynamic Bayesian network model that analyzes the continuous data of chromatin marks in 1-bp resolution for the entire genome. Both methods make *de novo* discoveries of major re-occuring patterns of chromatin marks.

ChromHMM and Segway were developed for analyzing a single epigenome. Although multiple epigenomes can be concatenated and hence analyzed together, such an approach ignores the critical information of position specificity of epigenetic events, and it treats all samples equally without accounting for cell type-specificity. Consequently, states generated by these methods are neither optimal nor robust for detecting epigenetic variation across multiple epigenomes. Extensions from ChromHMM and Segway have been developed to alleviate some of these issues. TreeHMM^18^ adds to ChromHMM a hierarchical model to borrow information across a known cell type hierarchy. A known cell type hierarchy, however, is not always available or informative, and a fixed hierarchy applied globally to the whole genome is overly restrictive. HiHMM^19^ extends from Segway to handle multiple epigenomes via infinite-state hidden Markov models (iHMM^20^). While each epigenome has its own iHMM parameters, information is shared across multiple epigenomes via priors. HiHMM however ignores the position specificity of epigenetic events.

We introduce IDEAS (Integrative and Differentiative Epigenome Annotation System), a new Bayesian framework for jointly analyzing multivariate epigenetic marks in multiple epigenomes. IDEAS is a powerful integration and segmentation tool that identifies *de novo* regulatory elements from high-throughput sequencing data in many samples. Simultaneously, IDEAS detects variation in the epigenetic patterns across both samples and genomic regions in varying lengths. Unlike existing approaches, we model cell type specificity locally across the genome. We also model position specificity via priors. We expect different cell types exhibiting similar signal patterns at the same loci, yet we allow variation in their posterior distributions. Combining local cell type and position effects, we leverage both global and local information from many samples to annotate regulatory elements and detect epigenetic variation more accurately than existing methods. We use Bayesian infinite-mixture models to approximate the quantitative data distributions rather than performing data binarization^16,21,22^ or making parametric distribution assumptions ^17,23-25^. While data binarization is sensitive to cutoff values and cannot detect quantitative variation, parametric assumptions on the data distribution are prone to model misspecification. Using mixture models, we are able to detect non-linear distributional variation beyond testing means in multivariate signals. We further utilize Bayesian non-parametric techniques to automatically determine the best model sizes (e.g., the number of states) without requiring technical inputs from the user. Our IDEAS is a unified framework that covers several existing tasks as special cases, such as genome segmentation, peak calling, differential gene expression testing. Importantly, IDEAS tackles these tasks in many samples simultaneously.

Using 84 ENCODE data sets in 6 cell types^26^, we demonstrate the superior accuracy and robustness of IDEAS for annotating regulatory elements and detecting epigenetic variation across cell types. The differential regulatory regions identified by IDEAS are driven by clustered cell type specific regulatory elements. They are strongly predictive of differential gene expression among cell types. By intersecting with disease variants from genome-wide association studies, we observed significant enrichment of these variants in cell type-specific regulatory regions, for which the phenotypes are highly relevant to the cell types. Current studies use specific epigenetic states enriched in certain cell types to perform disease enrichment analysis^5^. In comparison, we demonstrate that our modeling of the cell type-specific *differential* regulatory regions can yield much stronger enrichment scores with more interpretable phenotypes. IDEAS not only is a powerful tool for studying different gene regulation, but also provides an improved means for narrowing down the sets of plausible disease genetic variants and revealing cell type-specific connections between genetic variants and phenotypes.

## Results

### Overview of the Method

IDEAS is a Bayesian hierarchical mixture model that identifies multivariate patterns in high-throughput sequencing data and detects their variation across positions and samples in an unsupervised manner. The method is inspired by the haplotype inference problem for resequencing data. A haplotype inference procedure identifies combinatorial patterns of alleles shared among individuals over multiple loci. Similarly, our model identifies local clusters of cell types that share similar epigenetic landscapes over a region. In a haplotype inference problem, one needs to determine if a position is polymorphic or monomorphic. Similarly, our method assigns each position into a class summarizing its regulatory profile for all cell types, such as a plausible promoter, a potential enhancer, or a position with variable regulatory functions. A haplotype inference procedure calls specific alleles at each position in each individual, and its accuracy can be substantially improved by modeling haplotype structures in the region^27,28^. Similarly, our model assigns specific epigenetic states to each cell type at each position, with improved accuracy thanks to the incorporation of local cell type clustering and position classification. The output of IDEAS includes the inferred epigenetic states in each epigenome, local clustering of cell types, and genomic position classes, all obtained via automated Bayesian inference. An illustration of the IDEAS approach is shown in Figure 1.

**Figure 1.**
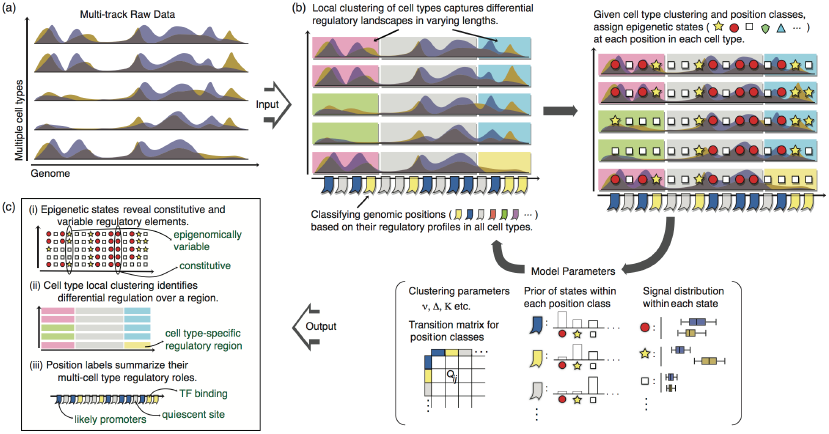
Illustration of the IDEAS model. (a) Input data include multiple tracks of epigenetic features in many cell types, possibly with replicates. (b) IDEAS fits the data iteratively by: 1) clustering cell types based on local epigenetic landscapes, where cell types carrying similar landscapes are clustered together; 2) assigning genomic positions into classes, such as potential enhancers, promoters, or insulators, using information in all cell types; 3) conditioning on cell type clustering and position classification, assigning an epigenetic state to each position in each cell type, where cell types clustered together at each position are more likely to be assigned the same epigenetic states; and 4) update model parameters. (c) Output of IDEAS includes the inferred epigenetic states like existing segmentation tools do. In addition, IDEAS outputs local cell type clusters that indicate potential differential regulation among cell types. IDEAS also reports position classes summarizing their regulatory profiles in multiple cell types.

### Annotation of Epigenetic States

We applied IDEAS to 84 ENCODE data sets in six human cell types (GM12878, H1-hESC, HeLa-S3, HepG2, HUVEC, K562), including 13 epigenetic features (the histone modifications H3K4me1, H3K4me2, H3K4me3, H3K9ac, H3K27ac, H3K27me3, H3K36me3, and H4K20me1, occupancy by POL2RA or CTCF, and chromatin accessibility monitored by Duke DNase, UW DNase, and FAIRE) and 1 control in each cell type. We took the maximum read counts in 200 bp nonoverlapping sliding windows, giving 13,763,197 windows genome-wide in each data set. Hoffman et al.^26^ have analyzed the same data using ChromHMM^16^ and Segway^17^. ChromHMM jointly segmented the six epigenomes via concatenation and generated 25 epigenetic states. Segway segmented each epigenome separately and generated 55 epigenetic states in total. We used the segmentation results of ChromHMM and Segway from Hoffman et al.^26^ for comparison, and we used similar mnemotics in Hoffman et al.^26^ to label the states by IDEAS.

#### Common and Novel States

IDEAS identified 24 epigenetic states (Figure 2a) in the six cell types. Overall, they agreed well with the states identified by ChromHMM (Supplementary Figure S1) and Segway. Several states were commonly identified, including transcription start sites, enhancers, CTCF occupancy, poised promoters, repressors, low and quiescent regions. IDEAS also identified some novel states. For examples, *TssCtcf* is a CTCF occupancy state highly enriched near transcription start sites, which has stronger active marks and open chromatin than the *Tss* state. *HistPol2* has much stronger Pol2 signals than the Pol2 state of ChromHMM. *HistPol2* was significantly enriched near genes involved in nucleosome organization (Binomial FDR 10^-56^), the *HIST* gene families (Binomial FDR 10^-53^) and many other core molecular processes (e.g., protein binding, gene translation). Remarkably, *HistPol2* occurs 10 times more frequently in the embryonic stem cell type (H1- hESC) (67% of *HistPol2,* Supplementary Figure S2) than in the other cell types (6%, 7%, 6%, 5%, 9% of *HistPol2* in GM12878, HeLa-S3, HepG2, HUVEC, K562, respectively). *BivProm* is a state with both active histone mark H3K4me3 and repressive histone mark H3K27me3. *BivProm* also has strong signals of the enhancer mark H3K427ac and is enriched near transcription start sites.

**Figure 2.**
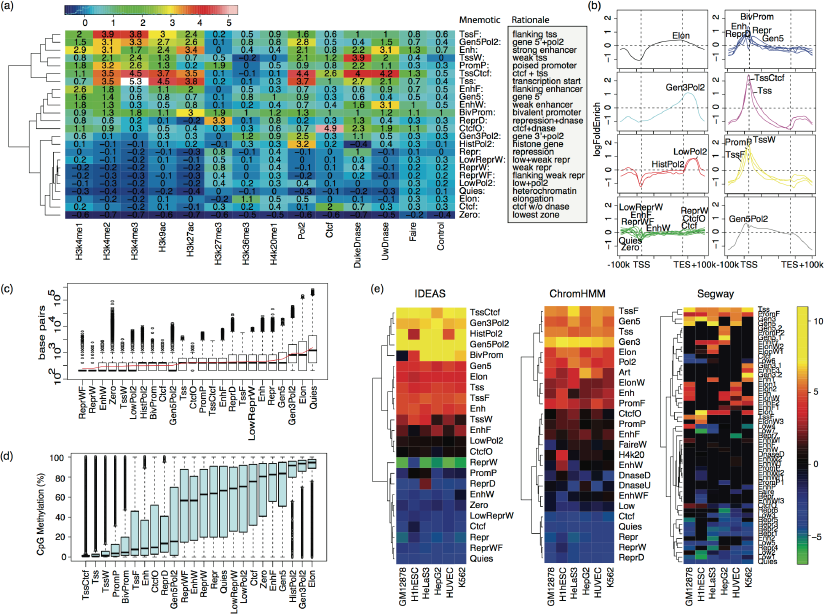
States generated by IDEAS. (a) Heatmap of the mean signals of the states inferred by IDEAS. State Mnemotics and a brief interpretation are shown on the right. A majority of the mnemotics is determined by overlapping the states of IDEAS with the states of ChromHMM and then checking the most overlapped states. (b) Log-fold enrichment of the states by IDEAS relative to transcription start sites (left dashed lines) and end sites (right dashed lines) clustered in 8 groups. (c) Boxplot of the lengths of IDEAS states. Red line indicates the mean lengths. (d) Boxplots of the percentage of CpG methylation in IDEAS states. (e) Heatmaps of the state effects on gene expression estimated by regressing the states within 2kb upstream of transcription start sites. RNA-seq data were normalized by the gene length followed by log2 transformation. The regression coefficients can be interpreted as the average increase in gene expression if a corresponding state is present within 2kb upstream of the TSS, conditioning on the other states. For genes whose transcript start sites were within 500bp to each other, as defined in GENECODEv7, only the left most gene was included.

IDEAS states exhibited unique spatial distributions (relative to the transcription start and end sites of genes in GENCODE^29^ v7) that corresponded well to their biological roles (Figure 2b). Tss-related states (*Tss, TssCtcf, TssW, TssF, PromP*) were enriched near transcription start sites and depleted near transcription end sites; repressors (*Repr* and *ReprD*), strong enhancers (*Enh*), bivalent promoters (*BivProm*) and Gene 5’ regions (*Gen5, Gen5Pol2*) were enriched in the flanking regions of transcription start sites; Elongation (*Elon*) and Pol2 states (*Gen3Pol2, LowPol2, HistPol2*) were depleted at transcription start sites but enriched near transcription end sites; low signal states (*Zero, Quies, ReprW, LowReprW, ReprWF*), weak marks (*EnhW* and *EnhF*), and *Ctcf, CtcfO* were neither obviously enriched nor depleted around genes. In addition, the lengths of IDEAS states varied considerably from 200bp to 100kb (Figure 2c). While most states were <1000bp on average, *LowReprW, Elon* and *Quies* had notably longer lengths.

#### CpG Methylation and Effects on Gene Expression

Overlapping IDEAS states with DNA methylation in all cell types revealed a clear bipartition of IDEAS states lying in either low or high methylation regions (Figure 2d). States carrying active epigenetic marks and centered around the transcription start sites (e.g., *TssCtcf, Tss, TssW*) have the lowest methylation levels, while states enriched at the transcription end sites (e.g., *HistPol2, Gen3Pol2, Elon*) have the highest methylation levels. These results are consistent with the substantial depletion of DNA methylation in active promoters and elevation in transcribed gene bodies^30^. Interestingly, among the states enriched in the flanking regions of TSS, e.g., *Repr, ReprD, Enh, Gen5,* the repressor states (*Repr, ReprD*) and *Gen5* are relatively highly methylated, but the strong enhancers (*Enh*) are not.

We further evaluated the state effects on gene expression. In each cell type, we fitted the RNA-Seq data of genes (GENCODEv7) by a linear regression model against the proportion of states within 2kb upstream of their transcription start sites. As expected, several of the states revealed by IDEAS had positive effects on gene expression, including the *Tss*-, *Pol2*- related states and *Enh, Elon, Gen5* and *Gen3* (Figure 2e). In contrast, repressors (*Repr, ReprD, ReprW*), insulators (*Ctcf*), *Quies* and *LowReprW* had negative effects on gene expression. Surprisingly, the three types of CTCF states (*TssCtcf*, *CtcfO*, *Ctcf*) identified by IDEAS showed opposite impacts on gene expression: *TssCtcf* has notably stronger positive effects than *Tss* on gene expression, suggesting that CTCF in the *TssCtcf* state facilitates transcription; *Ctcf* has strong negative effects on gene expression, suggesting an insulator role; and *CtcfO* has almost no effect on gene expression, which may indicate a non-regulating function of CTCF. The *ReprW* state resolved by IDEAS had a very strong negative effect on expression.

ChromHMM yielded similar state effects on gene expression, but the effects had smaller magnitudes and were less homogeneous across cell types. The state effects by Segway were much more heterogeneous across cell types due to the differences in state assignments in different cell types, and general trends across cell types were more difficult to detect. Overall, IDEAS produced the most homogenous state effect estimates across cell types, reflecting the benefit of including multivariate data from all cell types throughout the modeling.

#### Prediction of Enhancers

We evaluated the predicted enhancers using four independent validation datasets: 1) ENCODE EP300 peaks in GM12878, H1-hESC and HepG2 (note that EP300 was not included in the input data); 2) VISTA enhancer library^31^ containing 1699 experimentally tested loci with 882 verified enhancers; 3) 169 experimentally tested enhancers by FANTOM5^32^ in HeLa-S3 and HepG2; and 4) FANTOM5 CAGEtags that are predictive of active enhancers in GM12878, HeLa-S3, HepG2 and K562.

IDEAS captured 62% of EP300 peaks in its enhancer states (*Enh, EnhF, EnhW*) and 13% in *TssCtcf,* totaling 75% of EP300 peaks (Figure 3a). In comparison, ChromHMM captured only 41% of EP300 peaks in its enhancer states (*Enh, EnhW, EnhF*), and a significant proportion (33%) of EP300 peaks in its *Tss* state, totaling 74% of EP300 peaks. Segway captured 46% of EP300 peaks in its enhancer states (*Enh, Enh1, Enh2, EnhF1, EnhPr*), 17% in its *Tss* state, and 10% in *PromF* and *PromP2,* totaling 73% of EP300 peaks. Although all methods captured three quarters of EP300 peaks in a few states, IDEAS obtained the largest overlap with EP300 peaks in its enhancer states, and the fold enrichments of EP300 peaks were similar among the methods (Figure 3a). Further comparing the EP300 signals within the predicted strong enhancers (*Enh*) suggests that IDEAS strong enhancers tend to carry greater EP300 signals than the strong enhancers predicted by ChromHMM (Enh), Segway (*Enh/1/2*) and EnhancerFinder (Figure 3b).

**Figure 3.**
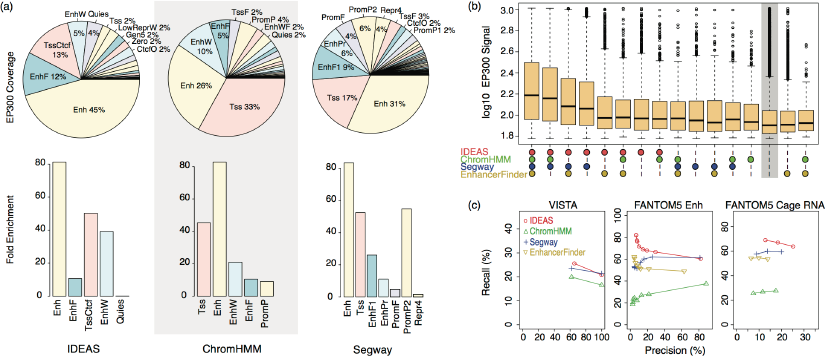
Comparison of predicted enhancers. (a) Percentage and fold enrichment of EP300 peaks captured by the inferred states (fold enrichment is shown only for states capturing >4% of EP300 peaks). (b) Boxplot of EP300 signals in the commonly and uniquely identified strong enhancer states. Each box shows the Log10 EP300 signals at the positions that are predicted as strong enhancers by certain combination of programs (marked by circles), and the grey area marks the positions not predicted as strong enhancers by all programs. (c) Precision and recall plots for the predicted strong enhancers, using VISTA enhancers, FANTOM5 enhancers, and FANTOM5 Cage RNA peaks as references, respectively. The plots were calculated using only the sites within each reference set, where positive and negative sites were determined using varying cutoff values. VISTA enhancers include validated and failed to validate enhancers in transgenic mice. We used two thresholds: one using the validated sites as true enhancers, and the other using all sites as true enhancers. For FANTOM5 enhancers, we used the experimental validation p-values as cutoffs. FANTOM5 called CAGE peaks as the predicted enhancers. We used the number of times (1, 2 or 3) a peak was called in 3 replicates as the cutoffs for true positives. EnhancerFinder was not compared in VISTA data because it was trained on VISTA enhancers.

We calculated the precision-recall rates (Figure 3c) comparing the strong enhancers predicted by IDEAS, ChromHMM, Segway and EnhancerFinder^33^ with the experimentally verified enhancers (VISTA and FANTOM5) and the CAGEtag data. IDEAS consistently achieved greater recall rates than ChromHMM and Segway in all validation data sets at different precision levels. IDEAS also performed better than EnhancerFinder on the FANTOM5 data sets. Since EnhancerFinder is a supervised learning method trained on VISTA enhancers, the stronger performance of IDEAS in finding FANTOM5 enhancers suggests that our unsupervised learning method may have a unique advantage in uncovering novel enhancers, particularly for those not represented in the training set.

#### Annotation of CTCF Binding

We evaluated the CTCF states using CpG methylation levels. The *TssCtcf, CtcfO* and *Ctcf* states by IDEAS showed distinct CpG methylation levels (Supplementary Figure S3). In comparison, the *Ctcf* and *CtcfO* states by ChromHMM and Segway had a mixed spectrum of methylation, with some *CtcfO* states showing much higher methylation levels than those in *Ctcf* states. We further compared the CTCF states with 51 validated CTCF binding sites in transgenic mice and 25971 putative CTCF sites from non-ENCODE experiments in CTCFBSDB2.0^34^. A CTCFBSDB2.0 site was counted as detected if the site was annotated as a CTCF state in any of the 6 cell types. IDEAS captured 48.1% of the validated CTCF sites and 92.3% of the putative CTCF sites (Supplementary Figure S3). In comparison, ChromHMM captured 21.2% validated and 73.9% putative sites, and Segway captured 25.5% validated and 78.0% putative sites. The fold enrichments of CTCF sites captured by the CTCF states of all methods were similar, indicating the best performance of IDEAS.

### Detection of Epigenetic Variation

We call a 200bp window epigenomically “variable” if the unit is assigned with different states in different cell types, otherwise the position is “constitutive". IDEAS marked 13.9% of the genome as variable sites. In contrast, 84.4% and 99.1% of the genome were marked as variable sites by ChromHMM and Segway, respectively. The substantially larger proportions of “variable sites” generated by ChromHMM and Segway may result from their inability to incorporate position specificity of epigenetic marks. Among the three methods, IDEAS generated the most homogeneous state assignments at each position in the six cell types (Figure 4a), and its cumulative state proportions were the most homogeneous across the six cell types (Supplementary Figure S2). Using the constitutive sites as a reference, analysis of variance suggested that the variable sites by IDEAS indeed carried substantial variability in the epigenetic features, whereas many of the “variable sites” by ChromHMM and Segway may actually be constitutive.

**Figure 4.**
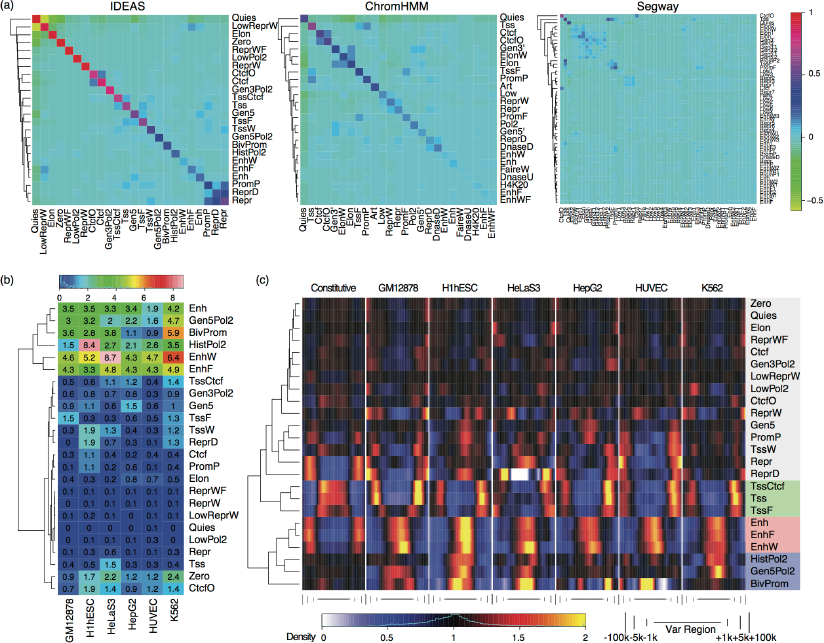
State co-occurrence and variation. (a) Correlation coefficients for states co-occurring at the same positions. (b) Heatmap of fold enrichment of states at cell type-specific sites relative to states at any variable sites non-specific to the cell type. (c) Density of states near and within constitutive and cell type-specific variable regions. For the cell type-specific variable regions, only states in the corresponding cell types are considered.

IDEAS was able to detect cell type specific states, such as enhancers (*Enh, EnhF, EnhW*) that had many fewer co-occurrence than the other states did (Figure 4a). There were many pairs of states that co-occur at the same positions significantly more frequently than by chance. *CtcfO* and *Ctcf*, for instance, substantially co-occurred at the same positions, as noted by IDEAS and ChromHMM. We also observed positive co-occurrence of Tss states (*Tss*, *TssF, TssW, TssCtcf*) and repressive states (*Repr, ReprD, PromP*). We noted a positive co-occurrence between strong enhancers (*Enh*) and repressed regions (*LowReprW*), which was revealed only in IDEAS (Figure 4a). For all cell types, the cell type specific variable sites were significantly enriched in enhancers (*Enh, EnhF, EnhW*). For some cell types, we also observed enrichment of *Gen5Pol2, HistPol2* and *BivProm* (Figure 4b). The cell type specific variable sites were significantly enriched near TSS (Supplementary Figure S5a), and the spatial distributions for different pairs of co-occurring states were distinct even after adjusting for the marginal spatial preference of each state in a pair (Supplementary Figure S5b).

We further detected long intervals showing consistently differential signals among cell types, which we call “variable regions”. Correspondingly, intervals showing similar signals in all cell types are called “constitutive regions”. IDEAS detected 70,622 variable regions of length >1kb, accounting for 26.5% of the genome. Among them, 69.3% were cell type specific (16.0%, 9.1%, 7.7%, 14.5%, 14.0% and 7.9% for GM12878, H1hESC, HeLaS3, HepG2, HUVEC and K562, respectively, covering 4.7%, 2.7%, 2.2%, 4.3%, 4.2% and 2.4% of the genome, respectively). Interestingly, a significant portion (75.8%) of the 200bp variable sites was located within the variable regions, suggesting that individual variable sites were locally clustered.

#### State Distribution Around Variable Regions

We observed a significant enrichment of enhancers (*Enh, EnhW*, *EnhF*) in the cell type specific variable regions, and they were depleted in constitutive regions (Figure 4c). Other states such as *HistPol2*, *Gen5Pol2* and *BivProm* were also enriched in the cell type specific variable regions, but only in some cell types. At those positions carrying enhancers, *Gen5, HistPol2* or *BivProm* in specific cell types, the corresponding states in the remaining cell types were mainly *Elon, LowReprW, Quies* (Supplementary Figure S6), with a few exceptions. For instance, several H1hESC-specific regions were defined by *HistPol2* and poised promoter (*PromP*), and the same positions were annotated as *Quies* and *Repr* in the other cell types, respectcely. Several HeLaS3-specific regions were repressed (*Repr*) but were poised promoters (*PromP*) in the other cell types. According to GREAT^35^, those regions were significantly enriched in targets of H3K27me3 and Polycomb proteins EED, Suz12, PRC2 that maintain transcriptionally repressive state of genes.

TSS states (*Tss*, *TssF, TssCtcf*) were significantly enriched in the close vicinity (<5kb) outside the boundaries of variable regions and within constitutive regions (Figure 4c). This suggests that variable regions tend to occur near genes and may influence differential gene expression. To test this hypothesis, we regressed gene expression data on the cell type partitions specified by each variable region. Using −log10 p-value of the regression model as weights, we computed a weighted frequency of TSS near or within all types of variable regions (but we skipped those types of variable regions with total length <5Mb). We observed a significant enrichment of TSS whose gene expression was associated with the variable regions (Supplementary Figure S7a). There were three TSS hotspots: within the variable regions and in their vicinities (1∼5kb) on both sides. The adjusted *r*^*2*^ for gene expression (Supplementary Figure S7b) explained by the variable regions was at the maximum for transcription start sites located within the variable regions, i.e., when a promoter and its gene body both carried differential epigenetic marks.

### Enrichment of Disease Variants

We overlapped the cell type specific variable sites and regions with the GWAS Catalog^36^ SNPs, including all lead SNPs and dbSNPs in strong LD with the lead SNPs (r^2^≥0.95). We calculated enrichment of disease variants relative to random shuffling of regions, and we identified 85 significantly enriched phenotypes. As shown in Figure 5a, our cell type specific variable sites and regions were significantly enriched in disease variants, and the enrichments were highly cell type specific. The enrichment patterns for the variable sites and the variable regions were similar. In fact, 98.6% of the cell type specific variable sites that contained at least one disease variant in Figure 5a were located within a cell type specific variable region. Only 1.4% of the disease enriched variable sites were singletons (relative to 24.2% by chance), suggesting that disease variants tend to occur in regions that demonstrate cell type specific activities over multiple positions.

**Figure 5.**
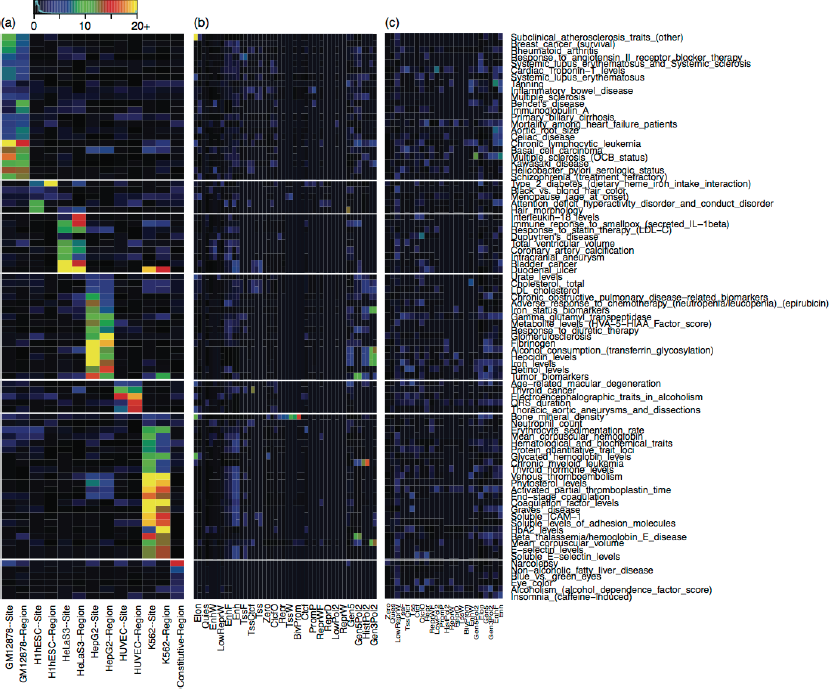
Disease enrichment in the detected variable regions. (a) Enrichment of disease variants in cell type-specific variable sites, variable regions and constitutive regions. GwasCatalog SNPs and dbSNPs in strong LD (*r*^2^≥0.95) with the lead SNPs were included. Only enrichments with Fold≥2, FDR≤0.05, and - log10(FDR)+Fold>=10 in at least one column in the matrix are shown. Enrichment and FDR were calculated by random shuffling of the variable sites and regions chromosome-wise. (b) Enrichment of disease variants in states that are unique to the most relevant cell types given in (a). There are two columns per state, for which enrichment was calculated against randomization within the cell type-specific sites and regions, respectively. (c) Enrichment of disease variants in states regardless of cell type specificity, for which randomization was done genome-wide in the most relevant cell types given in (a).

A majority of the phenotypes enriched in our variable sites/regions occurred in three cell types (GM12878, HepG2, K562), and they are highly relevant to the corresponding cell types. For instance, GM12878 is a lymphoblastoid cell line with a relatively normal karyotype. GM12878- specific sites are enriched in variants associated with several autoimmune diseases, including multiple sclerosis, rheumatoid arthritis, celiac disease and inflammatory bowl disease. HepG2 is derived from a liver carcinoma and is a model system to study metabolism disorders. HepG2-specific regions are significantly enriched in variants from metabolism related traits, such as iron levels, retinol levels, cholesterol levels and alcohol consumption. HepG2-specific regions are also enriched in variants for gamma-glutamyl transpeptidase and fibrinogen. While the former is most notably found in liver, the latter is mainly synthesized in liver. K562 is derived from a chronic myeloid leukemia (CML) patient, and K562-specific regions are significantly enriched in variants associated with CML and hemoglobin related traits. K562 has both erythoid and megakaryocytic properties, and appropriately, the K562-specific regions were enriched in variants associated with blood clotting, coagulation and erythroid phenotypes.

Our results suggest that identifying epigenetic cell type specificity lead to increased odds for finding genetic variants associated with cell type relevant phenotypes. In support of this, we found fewer and much weaker enrichment of disease variants in the constitutive regions (Figure 5a). While previous studies^5,12^ have used epigenetic states or specific epigenetic marks to narrow down regions for finding disease variants, our analysis shows that combining data from multiple cell types and identifying cell type specific differential regions may provide a more powerful means to find plausible disease variants. As shown in Figure 5b, within cell type specific variable sites and regions, disease variants were enriched in different types of epigenetic states for different phenotypes. Variants of K562-enriched phenotypes, for example, were mostly enriched in enhancers. Variants of HepG2-enriched phenotypes, on the other hand, tended to enrich in *Gen5Pol2, HistPol2* and *Gen3Pol2* regions. Although disease variants were overall more enriched in certain epigenetic states, the enrichments were generally in smaller magnitudes than using cell type specific variable regions (Figure 5c), and they did not suggest cell type relevance as our approach did. In addition, we performed the same enrichment analysis using ChromHMM and Segway segmentations. ChromHMM yielded fewer enriched phenotypes and many of them had no obvious connections to the corresponding cell types (Supplementary Figure S8). Segway resulted in just one enriched phenotype (aortic stiffness) without obvious relevance to GM12878.

## Discussion

While epigenetic states generated by segmentation tools can provide interpretable synopses for mapping the complex epigenetic landscapes, IDEAS extends the task to annotate multiple epigenomes via a unified probabilistic framework. By explicitly modeling position and cell type dependent events and leveraging information from local epigenetic landscapes in all samples, the inferred chromatin states are directly comparable across samples, allowing detection of both constitutive and differential regulatory events. IDEAS is more accurate for annotating regulatory elements than current state-of-the art algorithms, and simultaneously, it detects epigenetic variation across samples in regions of varying sizes. Our results revealed epigenetic cell type specificities and their impacts on differential gene expression across cell types. By intersecting the variable regions with non-coding SNPs from GWAS data sets, we observed a strong enrichment of genetic variants associated with human phenotypes, and the enriched phenotypes were highly relevant to the corresponding cell types. In an improvement over existing disease enrichment analysis, our modeling of the epigenetic cell type specificity yielded the strongest enrichment scores without constraining ourselves to specific epigenetic states. In comparison, disease enrichment analysis focusing on individual epigenetic marks or states, with or without considering cell type specificity, yielded smaller enrichment scores. While phenotypes often have cell type preferences, our results suggest that the regulatory mechanisms underlying a phenotype could be diverse. Our approach of directly addressing the epigenetic cell type specificity, therefore, provides a new tool for pinpointing plausible disease variants and interpreting their functions towards phenotypes in a cell type-specific context.

Additional applications of IDEAS include, but are not limited to, jointly calling peaks for constitutive and variable binding of transcription factors, testing differential gene expression in multiple conditions, and generalized association mapping of genetic, epigenetic and genomic data with phenotypes. For peak calling, IDEAS can quantify TF binding affinity using multiple states and detect co-binding of multiple transcription factors. For detecting differential gene expression, input data will be the expression level per gene normalized by the gene size, and differential expression can be reflected by different states. For generalized association mapping, diverse data types in different dimensions may first be integrated into categorical states by IDEAS, and then existing categorical testing procedures can be used to detect associations with phenotypes. The probability functions used in the mixture model of IDEAS can also be replaced by other choices so that it can be generalized to solve even a broader scope of problems.

This study attempts to integrate and characterize multiple epigenomes in both the dimensions of space (e.g., position along the genome) and time (e.g., stages of cellular differentiation). As high-throughput sequencing data sets are continuously generated for many more genome-wide features in additional cell/tissue types and conditions, large-scale computational methods like ours will be needed to facilitate in the analysis and interpretation of the ever growing dimensions of biological data, with a goal of untangling the complex mechanisms of gene regulation and revealing their contributions to phenotypes. While new insights generated from the results of IDEAS may be dependent on the data quality, the available epigenetic features and the cell types included, the initial run of our method on current ENCODE data is already useful for identifying DNA sequences in functional classes, finding regions of the epigenetic landscape that specific or selective for cell types, identifying key features driving the variation, and associating specific cell types with phenotypes of interest.

## URLs

ENCODE data sets analyzed in Hoffman et al. 2013 were downloaded from web portal: https://sites.google.com/site/anshulkundaje/projects/wiggler

ChromHMM and Segway segmentation results, HAIB Methyl RRBS DNA methylation data, Caltech RNA-seq data (we only used the first two replicates), EP300 peaks and raw signals, were downloaded from UCSC genome browser under ENCODE analysis hub: http://genome.ucsc.edu/ENCODE/analysis.html

VISTA enhancers were downloaded from VISTA enhancer browser: http://enhancer.lbl.gov/

FANTOM5 enhancers and CAGEtags were downloaded from FANTOM5 website: http://fantom.gsc.riken.jp/5/

EnhancerFinding predictions were downloaded from website: http://www.capralab.org/enhancerfinder-paper-published-in-plos-comp-bio/

Experimental CTCF sites and non-ENCODE putative CTCF sites were obtained from CTCFBSDB 2.0: http://insulatordb.uthsc.edu/homenew.php

GWAS Catalog and GENCODE files were downloaded from UCSC genome browser: http://genome.ucsc.edu/

## Acknowledgements

This work was supported by the National Institutes of Health grants R01DK065806 and R56DK065806 (YZ, RCH), R01HG004718 (YZ), RC2HG005573 and U54HG006998 (RCH). This work was supported in part through instrumentation funded by the National Science Foundation through grant OCI-0821527 (the Penn State CyberSTAR and BioSTAR computers).

## Author Contributions

Y.Z. and R.H. conceived of the project, Y.Z. designed the method and performed data analyses, and Y.Z., Y.F. and R.H. wrote the manuscript.

## Materials and Methods

### Data Preparation and IDEAS Specifics

The 84 ENCODE data sets in the 6 cell types and the segmentation results by ChromHMM and Segway were both obtained from Hoffman et al.^26^. The downloaded data sets have already been preprocessed and normalized. We followed the same procedure taken by ChromHMM to take the maximum read count per 200bp window as the input to our method, but we took log2(*x*+1) transformation of the read count (*x*) to reduce the skewness of read count distribution.

We used a hybrid of MCMC sampling and maximization steps to fit the IDEAS model. Starting from random values, we ran a mix of MCMC sampling and maximization iterations. The sampling steps were used for the purpose of jumping out of local modes of the likelihood surface. For the first 25 iterations, with probability *p* we ran maximization, and other wise we ran MCMC sampling, where *p* was increased from 0 to 0.5 linearly over the course of 25 iterations. For the next 25 iterations, we ran maximization and MCMC sampling with equal probability. After that, we ran additional 50 iterations of IDEAS by maximization only. The final output was taken as the results produced in the last iteration. In each iteration, we calculated the likelihood of the full model, which showed that the algorithm converged quickly after the 50^th^ iteration and stabilized at the end of the 100^th^ iteration. When running IDEAS, we further imposed a lower bound of 0.8 on the standard deviation of each epigenetic mark in each state, which was effective for smoothing out the discreteness of read count data, especially at low read count values. Consequently, we obtained fewer but much more interpretable states than otherwise would have been generated by artifacts in the data. We removed 1 outlying state generated by IDEAS, as it contained extreme values with only 10 instances genome-wide.

### The IDEAS Model

#### An Infinite Mixture Component Model

Let *X*={*x*_*ij*_}, for *i*=1,…*N* and *j*=1,…,*L*, denote the observation data in *N* cell types at *L* ordered genomic positions. Each *x*_*ij*_could be a *n*_*i*_ by *p* matrix, where *n*_*i*_ denotes the number of replicates for cell type *i* and *p* denotes the number of epigenetic features. The total number of samples therefore is 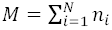. We model *X* by an infinite mixture of multivariate Gaussian distributions, where each Gaussian component has mean *μ*_*k*_ and covariance *Σ*_*k*_, for *k*=1,2,… ∞. We assign conjugate priors to *μ*_*k*_ and Σ_*k*_, *μ*_*k*_∼MVN(0, Σ_*k*_) and Σ_*k*_∼IW(*Φ*), where “MVN” stands for MultiVariate Normal distribution, “IW” stands for Inverse Wishart distribution, and *Φ* = *I*_*p*_. Let *K=*{*k*_*ij*_} denote the mixture component memberships of observations in *X*, where *k*_*ij*_=(*k*_*ij*1_,…,*k*_*ijni*_) includes memberships for *n*_*i*_ replicates. Let *m*_*k*_ denote the number of observations in the *k*th mixture component, and *X*_*k*_ denote the observations in the *k*th mixture component. It is standard^38^ to integrate out *μ*_*k*_ and Σ_*k*_ to obtain the following conditional distribution

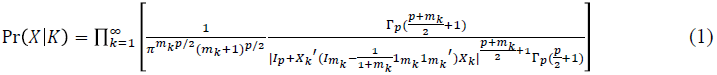

where Γ_*p*_(·) denotes a multivariate Gamma function. The term within the product in (1) is finite for finite samples, because the term equals 1 whenever *mk*=0.

#### A Hierarchical Mixture Model

An essential step is to model the distribution of mixture components *K*. Let *p*_*ij*_=(*p*_*ij*1_, *p*_*ij*2_,…) denote an infinite dimensional vector of proportions of mixture components for cell type *i* at position *j*. We treat {*p*_*ij*_} as random measures, i.e., *p*_*ij*_ is both cell type and position specific, but we impose constraints such that *p*_*ij*_ is not marginally independent across cell types and positions. We assume that cell types may belong to different groups at each position, where cell types in the same group at the position have the same random measure *p*_*ij*_, i.e., *p*_*ij*_=*p*_*i*’*j*_ if *i* and *i’* are in the same group at position *j.* This collapses the parameter space of {*p*_*ij*_} across samples. We further assume that different positions may belong to different classes, and each class has a specific prior distribution for the random measure *p*_*ij*_, i.e., *p*_*ij*_ and *p*_*ij*’_ follow the same prior distribution if *j* and *j’* belong to the same class. This imposes a hierarchy on the joint distribution of {*p*_*ij*_} across positions.

Let *S*={*s*_*ij*_} denote the group membership of cell type *i* at position *j,* and let *O*={*o*_*j*_} denote the class of position *j.* Given *s*_*ij*_=*s* and *o*_*j*_=*o*, we rewrite *p*_*ij*_ as *p*_*ij*_ = *f*_*s,o,j*_, where *f*_*s,o,j*_ is again an infinite dimensional vector for the proportions of mixture components, but it depends on sample group *s*, position class *o* and position *j*. Using this new parameterization, we borrow information across both samples and positions: cell types in the same group (*s*) at each position will yield same distribution of epigenetic features, and positions in the same class (*o*) will yield similar but not identical distribution of epigenetic features. Here, *S* serves as local clustering of cell types, and *O* provides summary information of genome positions for all samples.

We treat *f*_*s,o,j*_ as conditionally independent random measures given (*s*, *o, j*), and we model *f*_*s,o,j*_ by a Dirichlet Process^39^ with hyper parameter *Aα*_0_. The concentration parameter *A* (default = *N*) is a constant that balances between position-specificity (when *A* is small) and class-congruity (when *A* is large) for the distribution of mixture proportions. The proportion parameter *α*_0_ is specific to each position class *o,* and is an infinite dimensional vector taking values between [0,1] and sums to 1. Let *α* = {*α*_0_}, we have the conditional probability function for *K* and {*f*_*s,o,j*_} as

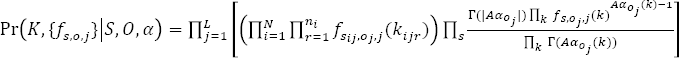

where operator |·| denotes the sum of all elements in the vector. Let {*c*_*sj*_(*k*)}denote the count of mixture component *k* in group *s* at position *j.* We integrate out {*f*_*s,o,j*_} to obtain the following conditional probability function

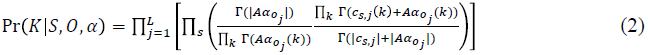

Again, the product over component *k* in (2) is finite for finite samples, as the term within the product is 1 for those *k* with *c*_*sk*_(*k*) = 0.

#### Local Clustering of Cell Types

We utilize iHMMs^20^ to model the local clustering variable *S* of samples across positions. The Markov property allows local dependence of clustering, and simultaneously, the infinite state model allows the number of clusters to be learned from the data. We model each cell type by one iHMM, *S*_*i*_=(*s*_*i*1_,…,*s*_*iL*_) for each cell type *i*, and *S=*(*S*_1_,…,*S*_*N*_)={*s*_*ij*_}. We assume independence between Markov chains, and all chains are governed by two sets of parameters: an infinite vector of state probabilities, and a *L*-dim vector of position-specific transition parameters.

Let *v=*(*v*_*1*_, *v*_*2*_,…) denote an infinite vector of state probabilities that sum to 1. We use a stick-breaking process^40^ to describe the prior distribution of *v.* Let (*V*_*1*_, *V*_*2*_,…) denote an infinite vector of independent *Beta* random variables, *V*_*s*_∼*Beta*(1,1) for s=1,2,…. We calculate *v* by *v*_*s*_=*V*_*s*_Π_*t*<*s*_(1- *V*_*t*_) for *s*=1,2,…. The posterior distribution of *v* is then a stick-breaking process. To model transition between states across positions, we utilize a simple mechanism: at each position *j*, the iHMM decides whether or not to select a new state; if yes, a new state is randomly selected from distribution *v,* otherwise the state at position *j* remains as the state at position *j*-1. Let {*r*_*j*_} denote a *L*-dim vector of transition parameters at *j=1,…*,*L*, with *r*_*1*_=1. Let *Δ*={*δ*_*ij*_} indicate whether or not a transition of state occurred in cell type *i* at position *j.* We model *δ*_*ij*_∼Bernoulli(*r*_*j*_) independently. The model for (*S*_*i*_, *Δ*_*i*_) for each cell type *i* is therefore

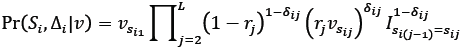

where the indicator 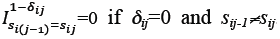. We do not infer {*r*_*j*_} but assign a prior *r*_*j*_∼Beta(**γ,**1-**γ_*j*_**)and integrate {*r*_*j*_} out, where 0<1*γ*_*j*_< 1 denotes a pre-determined constant (by default **γ**_*j*_= 1 — *e*^-0.1*d*_*j*_^ where *d*_*j*_ denotes the distance between positions *j* and (*j-*1) in Kb). Let 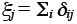 denote the total number of transition events at position *j* in all iHMMs, we obtain

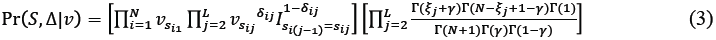

#### Classification of Genomic Positions

It is also natural to use iHMM to model the position class variable *0=*{*o*_*i*_,…,*o*_*L*_}. Let *q=*{*q*_*00*_,} denote an infinite square matrix of state transition probabilities such that each row sums to 1. We use a stick-breaking process to model the prior distribution of each row of {*q*_*00*_,}*-* For each row *o* in *q,* let {*U*_*0*_,}denote an infinite set of independent *Beta* random variables, *U*_0_,*∼Beta*(1,1), then the priori distribution of *q*_*00*_, is determined by 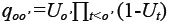. This prior does not favor a Markov chain to remain in the same states across positions, which could be unrealistic. As a remedy, we introduce *w =* {*w*_*o*_} denoting the probability that, given current state o, the state at the next position will be selected anew, and with 1–*w*_*o*_ probability the state at the next position will remain in *o.* We treat *w*_*0*_ as random with *w*_*0*_ ∼Beta(0.5,0.5). As a result, our transition probability matrix is designed as *q\** = diag(1 -*w*_*o*_) + *w*_*o*_*q*. Finally, we let the initial probability *p=* {*p*_*0*_}to be modeled by another stick breaking process using Beta(l,l). The probability function of *O* given parameters (*p, w, q*) can therefore be written as

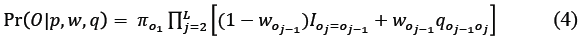

Combining formulas (l)-(4), we obtain the probability model:

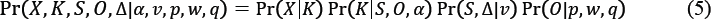

The parameters *α,v,p,w,q* in (5) are continuous variables, which we further assign prior distributions as follows: *α, v,p*, *q* consist of infinite dimensional vectors of proportions, each of which is modeled by a stick-breaking process prior via Beta(l,l), and *w* consists of Beta random variables with prior Beta(0.5,0.5) for each position class *o.*

###### Here is a summary of notations in the IDEAS model

<u>Constants:</u>

*N:* number of individuals;

*p:* number of features;

*L:* number of genomic positions;

*M:* total number of samples, which equals to the summation of all replicates;

*X*: *M*×(p*L*) data matrix of *p* features from *N* individuals at *L* positions;

*A:* concentration parameter for the prior of mixture proportions;

*γ:* position-specific parameters for the change-of-state probability of individual partitions;

<u>Discrete Variables:</u>

*K*: *M*×*L* matrix of mixture component memberships;

*S*: *N*×*L* matrix of individual partitions;

*Δ*: *N*×*L* indicator matrix of transition events in individual partition *S;*

*O:* L-dim vector of position classes;

<u>Continuous Variables:</u>

*α:* collection of infinite-dim vectors of concentration parameters for the mixture component proportions in each position state *o;*

*v:* infinite-dim vector of individual state proportions;

*p:* infinite-dim vector of initial distribution of position states;

*w:* infinite-dim vector of the probabilities of state transition from position states *o;*

*q:* infinite-dim matrix of position state transition probabilities, if a transition event occurs;

## Model Inference Procedure A Hybrid of Maximization and MCMC

While standard MCMC algorithms are available to infer a Bayesian model, it may take indefinite time for the model to converge on genome-scale data sets. Since our main interest is to identify a plausible model that best describe the data, we need a faster inference procedure such as model maximization. In a complex model like ours, however, maximization may be easily trapped in local modes. We therefore implemented a hybrid between the two. In the first few iterations, we use MCMC to sample model parameters stochastically to explore plausible model structures. As the iteration moves along, we begin to maximize the model parameters with increasing frequency. Next, we describe our model fitting procedure in a sampling scheme, whereas maximization can be done by simply taking model maximizers.

### Update Mixture Membership (*K*)

We update *K* from Pr(*X*|*K*} Pr(*K*|*S, 0, α*) for one observation at a time using a Gibbs sampler. To update the *k*_*ij*_ for cell type *i* at position *j*, let 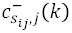 denote the number of observations in mixture component *k* in group *s*_*ij*_ at position 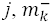 denote the total number of observations in component *k,* 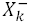 denote the collection of observations in component *k,* and 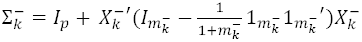 denote the unscaled covariance matrix (not standardized by sample size) for component *k*, all less the current observation to be updated for. Using (1) and (2), the probability for the current observation to be assigned to component *k* is given by

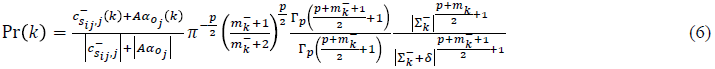

The first ratio term is from (2), the remaining terms are from (1), and δ denotes the change in the unscaled covariance matrix after adding the observation into component k. When 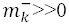, we can simplify the last ratio term to

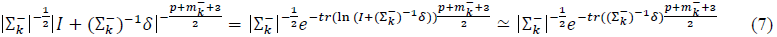

where the approximation is due to first order Tayler expansion. It can be shown that 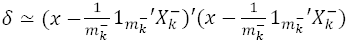, which is the square of the current observation (*x*) subtracting the estimated means. More details of approximation (7) can be found in *Supplementary Notes.*

We calculate the probability of assigning an observation to component *k* using (6). But to improve computing speed, we replace the last term in (6) by (7). Also, we practically replace 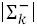 by |Σ_*k*_ |, which needs to be computed only once per iteration. Note that for any new component 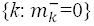, (6) is proportional to the hyper parameter 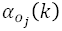. Consequently, we can collapse the infinitely many new components and calculate an overall probability of assigning the observation to a new component, and a new component is chosen according to 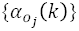.

### Update Cell Type Partition (*S*, *Δ*)

Given (*K,O,* α,*v*), we update (*S*, Δ) from Pr(*K*|*S, 0,* α) Pr(*S*, Δ|*v*). We update each iHMM *S*_*i*_ (and Δ_*i*_) conditioning on all other iHMMs. It is easily checked that the initial distribution of *S*_*i*_ is the infinite-dim vector *v*, the transition matrix is an infinite-dim matrix diag(1-*r*_*j*_)+*r*_*j*_1*v’*, where 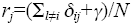 and the emission probability of mixture component *k* from state *s* at position *j* is 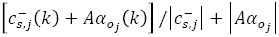 (this is for the simple case without replicates, i.e., *n*_*i*_=1, but the general case with replicates easily follows). To reduce the infinite number of states into a finite set, we notice that the emission probability is the same for all states *s* that do not appear in the remaining *N*-1 iHMMs, because 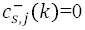 for *k*. Let *s\**=max(*S*_-*i*_) denote the maximum index of states in all other chains in *S.* We can temporarily reduce the state space of *S*_*i*_ to *s*\*+1 states, with the last state denoting a collapsed new state. Let *v*\* *=* (*v*_1_,…, *v*_*s*_*, *v*_*new*_)′, then *v*\* is a finite-dim vector denoting the initial state distribution of the reduced (finite) Markov model, and its transition matrix is also finite in form of diag(1-*r*_*j*_)+*r*_*j*_1*v* *’. We use a standard forward-backward sampling algorithm to update *S*_*i*_ and its transition indicator Δ_*j*_. In the forward step, we recursively calculate the joint probability function *Pr*(*k*_*i*1_,…*k*_*ij*_, *s*_*ij*_|*O,a,v\**) at position *j,* for *j=*1,2,…,*L,* by integrating out all possible paths in *S*_*i*_ and its transition indicator Δ_*i*_ up to position *j.* In the backward step, we sample *S*_*i*_ and Δ_*i*_, in the reverse direction for *j*=*L,L*-1,…,1, conditioning on the already sampled path (*s*_*ij*+1_,…,*s*_*iL*_) and Pr(*k*_*i*1_,…,*k*_*ij*_, *s*_*ij*_|*O,α,v*\*). Finally, for any positions assigned to the collapsed new state, we expand the state space back to the original infinite dimensions and assign each of those positions to a new state with index >*s*\* according to *v.* More details on collapsing the infinite states can be found in *Supplementary Notes.*

### Update Position Class (*O*)

Given (*K,S, α,p,w,q*), we update position class *O* from *Pr*(*K* |S, *0, α*) Pr(0|p, *w, q*). Again, we handle infinite dimensions by collapsing the empty states in *O* into one state so that the computation becomes finite. We use a standard forward-backward algorithm to update *O* in the collapsed Markov chain. We then expand the collapsed states back to the original infinite dimensions according to the priors specified by *p, w* and *q*. Collapsing is possible because the emission probability function of mixture components from an empty state *o* is specified by DirichletProc(A*α*_*o*_), and *α*_*o*_ for all unoccupied states *o* are identical.

### Update (*p,w,q*)

Assuming stationary Markov chain, the initial distribution *p* can be updated by first calculating 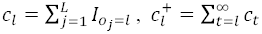, for *l*=1,2,…, then sampling 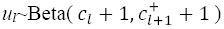, and computing 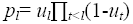. The transition event probability *w* is updated by counting the number *h*_*l*_ of transition events occurred in each state *l=* 1,2,…, and update *w*_*l*_∼Beta(0.5+*h*_*l*_, 0.5+*n*_*r*_*h*_*l*_). The transition matrix *q* is updated within each row separately using the same method for updating *p*, except that only those positions whose previous position is in the state of the corresponding row in *q* are considered. In any of these updates, we only update the first *o*_*max*_ rows and columns, where *o*_*max*_ denotes the maximum state index in the current vector *O*. For the remaining infinitely many elements in *p*,*w*,*q*, which correspond to the states that do not occur in the current position classes, we simply use their mean values as determined by the prior distributions.

### Update *α*

Condition on (*K, S, O*), we update *α*_*o*_ for *o*=1,2,…, where |*α*_*o*_|=1. We update α_*o*_ by counting the number of mixture component *k* within position state *o*, plus the prior parameter *Aα*_*o*_ using the current value of *α*_*o*_, and denote the total by *n*_*o*_(*k*); calculate 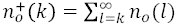 and generate random values *u*_*k*_ from Beta 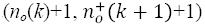; and then update 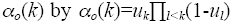, for *o*=1,2,…,max(*O*). For the remaining empty states, we use their mean values as specified by the prior distribution.

### Update *v*

Similarly, we use (*S*, *Δ*) to update *v.* We first calculate the number (*t*_*s*_) of transition events in all chains in *S* that move to state *s*, including the initial state *s*_*i*1_, *i*=1,…,*N*. We compute 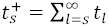 and generate random values *u*_*s*_ from Beta 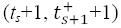; and then update *v*_*s*_ by 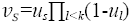, for *s*=1,2,…,max(*S*). For the remaining empty states, we again use their mean values as specified by the prior distribution.

## Supplementary Notes for Bayesian Modeling of Epigenetic Variation in Multiple Human Cell Types

### Gaussian Emission Probability

In our Bayesian model, we derive the emission density function from following joint distribution of Gaussian data *X*_*d*_ in component *d* with covariate *Z*_*d*_:

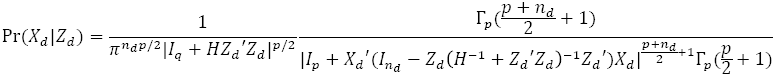

where *X*_*d*_ is a *n*_*d*_х*p* matrix, *Z*_*d*_ is a *n*_*d*_х*q* matrix, *n*_*d*_ denotes sample size, and *H* denotes a *q*х*q* hyper-parameter matrix.

Suppose a new data *x* (and covariate *z*) is added to component *d*. Let 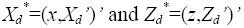. The conditional density is

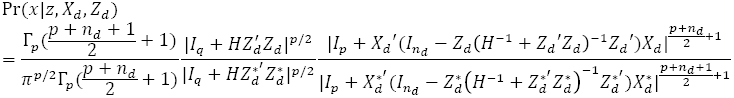

We first simplify

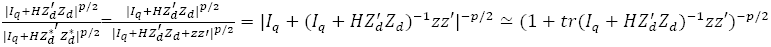

Next, let 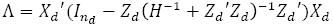 and 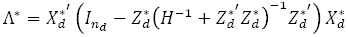, then

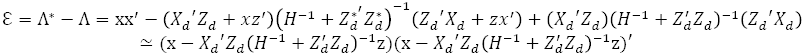

(This approximation assumes large *n*_*d*_)

Therefore 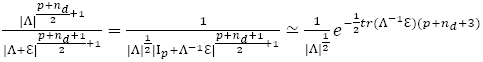

As a result, 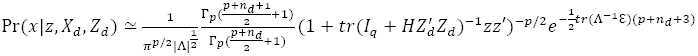.

### Forward Summation and Backward Sampling for Updating (*S*, *Δ*)

For each individual i, we fit a Markov chain and we use a forward summation backward sampling algorithm to update its variables (*Si, Δi*) conditioning on the parameters of the remaining individuals. For notation simplicity, hereafter we drop the individual index *i*. For *j*=1,…,*L* in ascending order, the forward-summation step sequentially computes the marginalized probability of the Markov chain starting from position 1 and ending at position *j,* with specific state *S*_*j*_ at position j. Reversely, the backward sampling algorithm sequentially samples a path of states staring from position *L* and ending at position *j,* for *j=L,…,1* in descending order.

Let *s*\* denote the maximum state index taken by all individuals in the current iteration. In the forward-summation step, we collapse all states whose indices >*s*\* into a super state, and we denote the super state by index *s*\*+1. Let *ssj* denote the state at position *j* after collapsing, then *ss*_*j*_ takes *s*\*+1 possible values. Further let *vv*_*s*_ = *v*_*s*_ for *s≤s\** and 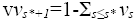 denote the probability of the super states *s*\*+1. The forward step computes the marginal probability at position 1 as

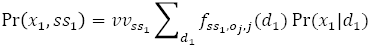

Here 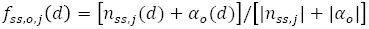, where *n*_*ss,j*_(*d*) denote the number of Gaussian component *d* in the remaining individuals in state *ss* at position *j*. Pr(*x*|*d*) denotes a Gaussian density function for component *d*, with 1^st^ moment 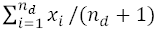 and 2^nd^ moment 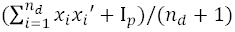.

Sequentially, we compute the joint probability of data at positions 1,…,j and simultaneously the Markov chain visit state *ss*_*j*_ at position *j* as

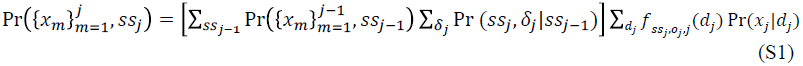

Formula (S1) first sums over all possible transitions from previous state at position *j*-1 to current state at position *j,* and then calculates the probability of emitting data *d*_*j*_ from the state *ss*_*j*_.

In the backward sampling step, we first sample the super states *ss*_*L*_ from the marginal probability at position L. If *ss*_*L*_=*s*\*+1, it means that the Markov chain visits a new state. In this case, we further update *s*_*L*_by *s*\*+*y*, where *y* denotes a random positive integer generated from *Geometric*(1*/*(1*+A*)), where *A* is the hyper-parameter we used in the Beta prior. This Geometric distribution is a result of our stick-breaking prior, as no data has yet visited this state. On the other hand, if *ss*_*L*_*≤s*\*, then *s*_*L*_=*ss*_*L*_. Given the states at position *j,* we sample the states at position *j*-1 sequentially in reverse order. We first sample the transition event indicators *δ*_*j*_ using the probabilities in formula (S1) (where the last summation term can be ignored), conditioning on the state at position *j.* We then sample the state at position *j*-1 according to *δ*_*j*_. Particularly, if no change of state at position *j*, then *ss*_*j*-1_=*ss*_*j*_ and *s*_*j*-1_=*s*_*j*_. Otherwise, we sample *ss*_*j*-1_ and *s*_*j*-1_ in similar ways as described for position *L*. Finally, given the newly generated states, we update the Gaussian component memberships {*d*_*j*_}.

## Supplementary Figure Legends

**Figure S1.**
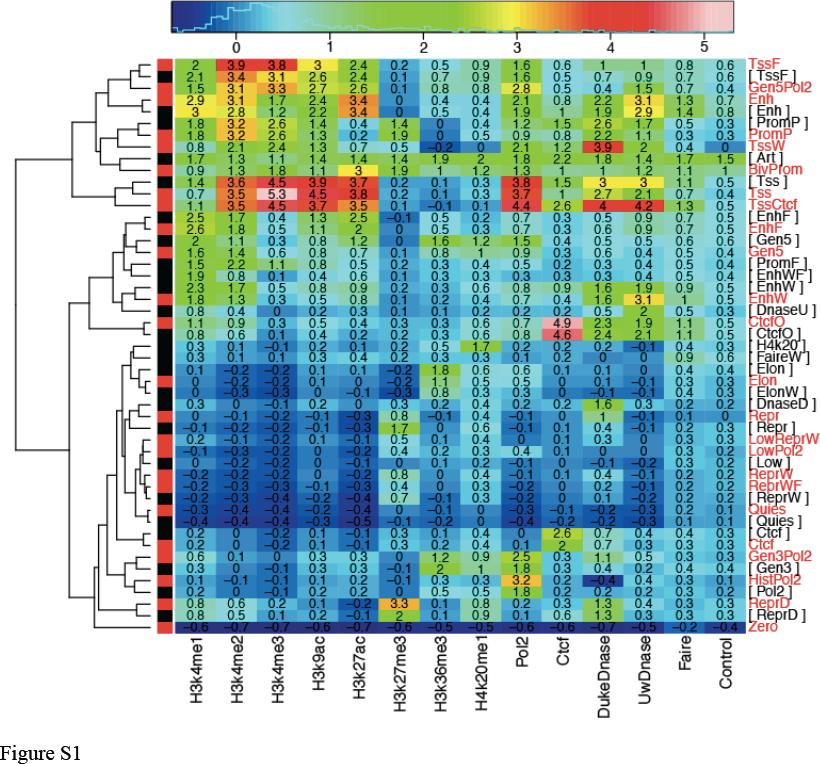
Heatmap of the mean signals of the states inferred by IDEAS (red labels) compared with the mean signals of the states inferred by ChromHMM (black labels in brackets).

**Figure S2.**
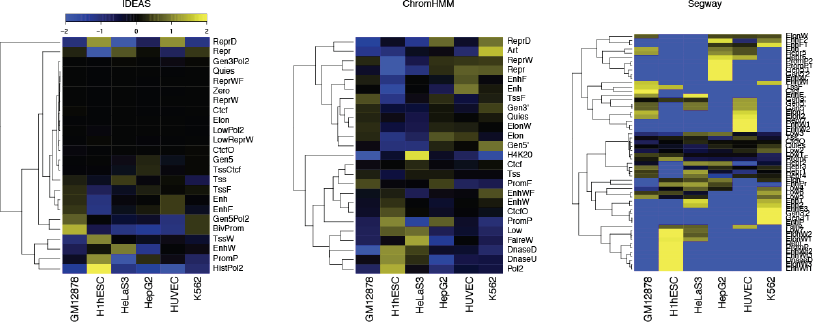
Log enrichment and depletion of states in the six cell types relative to equal distributions.

**Figure S3.**
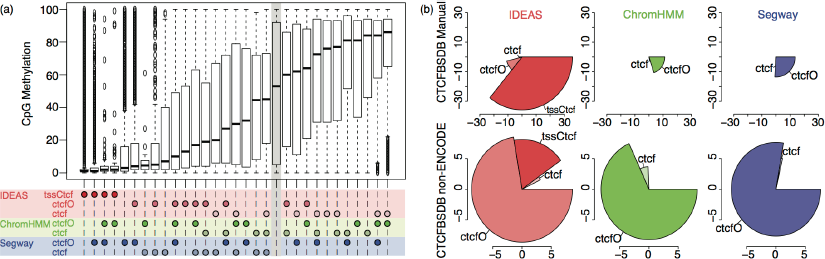
Comparison of predicted CTCF occupancy. (a) Boxplot of percentage of CpG methylation in the commonly and uniquely identified CTCF states. Each box shows the methylation levels at a set of positions that are predicted as CTCF by certain combination of programs (marked by circles), and the grey area marks the positions not predicted as CTCF by all programs. (b) Pie plots showing the power (angle) and the fold enrichment (radius) of each method using CTCFBSDB2.0 CTCF sites as references. The pies start from 3’ clock position and span in clockwise direction. Radius of a pie denotes the magnitude of fold enrichment of reference CTCF sites. Each pie corresponds to a CTCF state, and the angle of a new pie, after an existing pie, represents the extra proportion of reference CTCF sites captured by the corresponding CTCF state.

**Figure S4.**
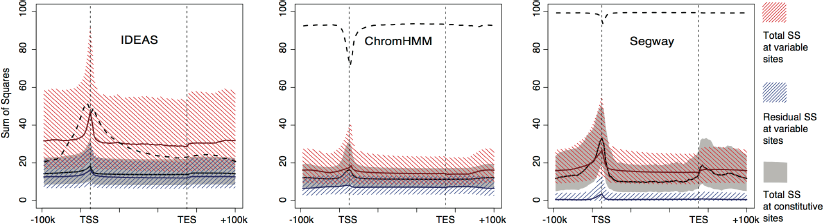
25^th^-75^th^ percentiles of the total and residual sum of squares (S.S.) of epigenetic signals at positions relative to TSS and TES. Red area shows the total S.S. at the detected variable sites; blue area shows the within-state S.S. at the variable sites; grey area shows the total S.S. at the constitutive sites. Note that the residual S.S. by ChromHMM and Segway is notably lower than the S.S. at their constitutive sites, suggesting that many of their “variable sites” may be attributable to noise. Medians are shown in solid lines. Percentage of variable sites relative to the total number of sites at each position is shown in dashed lines.

**Figure S5.**
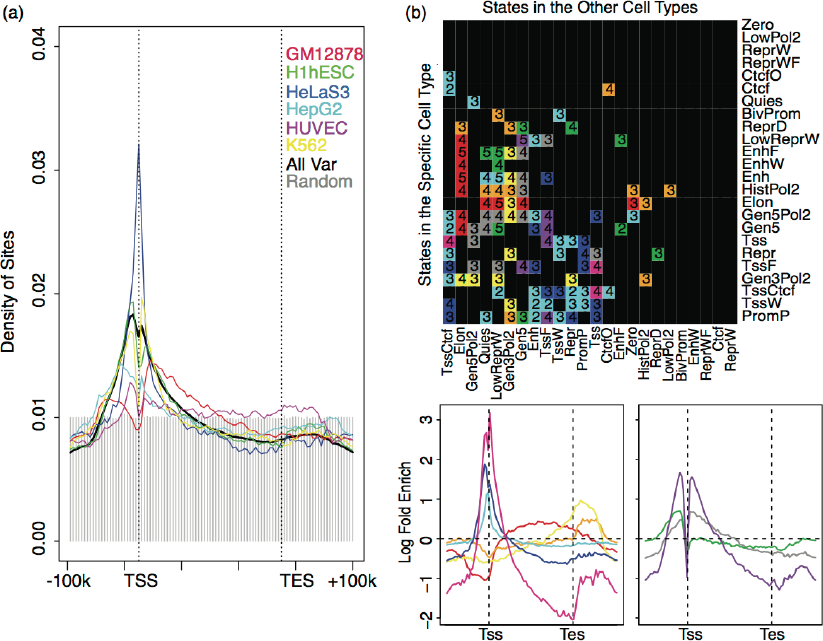
(a) Spatial distribution of cell type-specific variable sites relative to TSS and TES. Combined distribution of all variable sites is shown in black, and distribution of random sites is shown in grey area as a reference. (b) Spatial distribution of state pairs co-occurring at cell type-specific variable sites. Top panel is a color-coding matrix for the spatial distribution of each pair of states (black color means uniform distribution). Each row in the matrix corresponds to the state observed in the specific cell type, and each column corresponds to the state observed in the remaining cell type. Numbers shown in the matrix cells denote the log10 total count of each state pairs observed. Bottom panel shows the spatial log fold enrichment of the state pairs color-coded in the matrix above it. The enrichment was calculated relative to the product (independence) of the marginal spatial distributions of the states in a pair.

**Figure S6.**
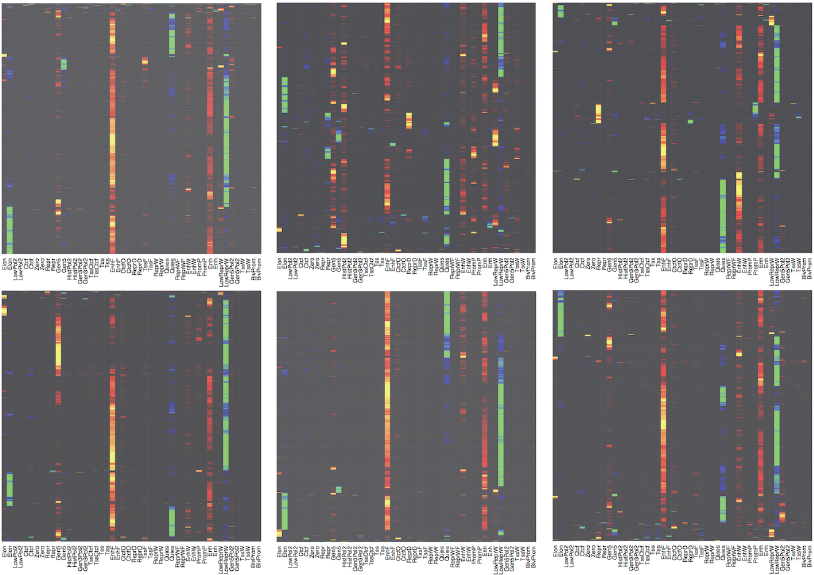
Distribution of states in cell type specific variable regions. Each row in the heatmap corresponds to a variable region. Warm colors (black to red to yellow for 0 to 0.5 to 1) indicate states observed in the corresponding cell types. Cold colors (black to blue to green for 0 to 0.5 to 1) indicate states observed in the other cell types. Only states in cell type specific variable sites are counted. Top row from left to right: Gm12878, H1hesc, HelaS3. Bottom row from left to right: HepG2, HUVEC, K562.

**Figure S7.**
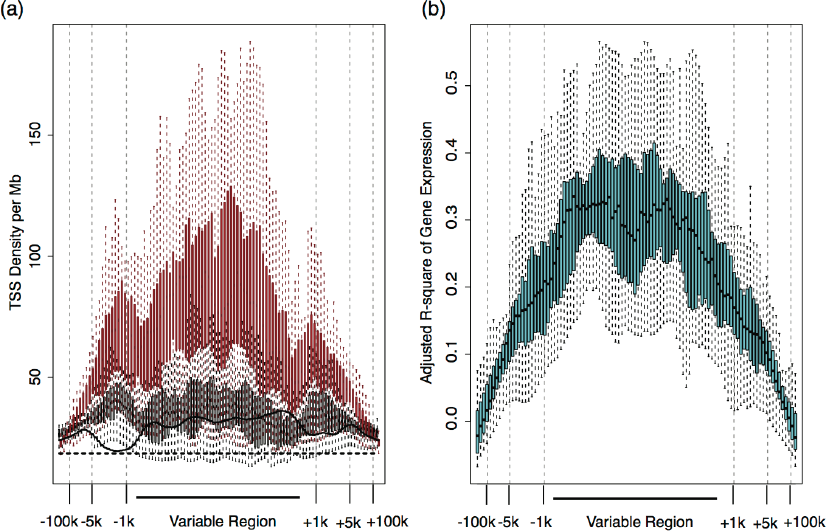
(a) Boxplots of TSS density (per Mb) near and within constitutive and variable regions. Red boxes show the TSS density weighted by −log10 p-value of association between gene expression and the variable region, black boxes show the unweighted TSS density, black line shows the TSS density around constitutive regions and dashed horizontal line shows the genome-wide average TSS density. (b) Boxplots of adjusted *r*^*2*^ for gene expression explained by the cell type partitions in the variable region.

**Figure S8.**
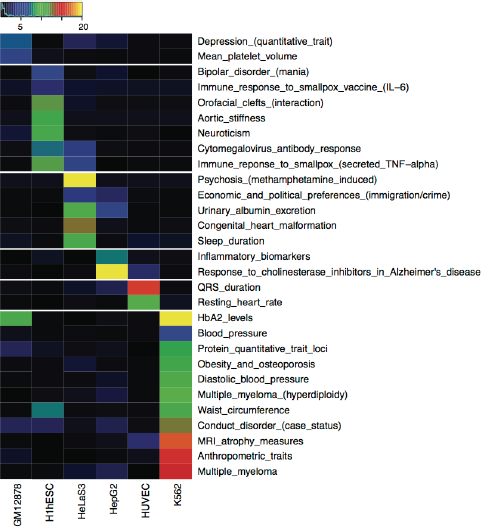
Enrichment of disease variants in ChromHMM cell type specific sites.

